# Regulator of calcineurin-2 is a centriolar protein with a role in cilia length control

**DOI:** 10.1101/188946

**Authors:** Nicola L. Stevenson, Dylan J.M. Bergen, Amadeus Xu, Emily Wyatt, Freya Henry, Janine McCaughey, Laura Vuolo, Chrissy L. Hammond, David J. Stephens

**Author notes:** These authors contributed equally to this work (Nicola L. Stevenson, Dylan J.M. Bergen).

## Abstract

Almost every cell in the human body extends a primary cilium. Defective cilia function leads to a set of disorders known as ciliopathies characterised by debilitating developmental defects affecting many tissues. Here we report a new role for regulator of calcineurin 2, RCAN2, in primary cilia function. It localises to centrioles and the basal body and is required to maintain normal cilia length. RCAN2 was identified as the most strongly upregulated gene from a comparative RNAseq analysis of cells in which expression of the Golgi matrix protein giantin had been abolished by gene editing. In contrast to previous work where we showed that depletion of giantin by RNAi results in defects in ciliogenesis and in cilia length control, giantin knockout cells generate normal cilia on serum withdrawal. Furthermore, giantin knockout zebrafish show increased expression of RCAN2. Importantly, suppression of RCAN2 expression in giantin knockout cells results in the same defects in cilia length control seen on RNAi of giantin itself. Together these data define RCAN2 as a regulator of cilia function that can compensate for loss of giantin function.

## Introduction

Ciliogenesis, the emergence of a microtubule axoneme from the mother centriole, is fundamental for the ability of non-cycling cells to sense and respond to their environment (Lechtreck, 2015). Defects in the formation and/or function of the cilium lead to a cohort of human diseases known as ciliopathies (Braun and Hildebrandt, 2016) which affect multiple tissues leading to problems with respiratory, kidney, and heart function as well as vision, hearing, and fertility. Cilia defects are also linked to obesity and diabetes. Movement of cargo and signalling complexes into and out of cilia requires intraflagellar transport (IFT), the process by which microtubule motors drive motility along the axoneme. The ability of cilia to act as confined signalling hubs directing key developmental and homeostatic pathways, such as those linked to sonic hedgehog (Shh) signalling (He et al., 2016) or mechanosensing, requires the trafficking of receptors and associated signalling molecules into cilia (He et al., 2016; Sung and Leroux, 2013). This includes ligand-gated receptors, G-protein coupled receptors, Ca^2+^ channels, and transcription factors. Considerable work supports a major role for Ca^2+^ in ciliary function (Delling et al., 2013). Resting cilium [Ca^2+^] is substantially higher than resting cytoplasmic [Ca^2+^] and ciliary [Ca^2+^] is important for cells to respond to Shh through activation of Gli transcription factors (Delling et al., 2013).

We have shown previously using RNAi that the golgin giantin is required for ciliogenesis *in vitro* (Asante et al., 2013). This was linked to the accumulation of the dynein-2 motor complex around the ciliary base. Dynein-2 is the major motor driving retrograde IFT along the axoneme and is, among other functions, required for the transduction of the Shh signal (He et al., 2016). Consistent with our findings using RNAi in cultured cells, morpholino knockdown of giantin in zebrafish resulted in fewer but longer cilia in the neural tube (Bergen et al., 2017). In contrast, recent characterisation of giantin KO zebrafish shows that, while breeding and developing grossly similar to wild-type fish, giantin knockout (KO) fish show a significant decrease in body length (more notable in young adults) and have defects in extracellular matrix, cartilage and bone formation, but only minor cilia defects (Bergen et al., 2017).

We recently generated a KO cell line that no longer expresses giantin (Stevenson et al., 2017). Here we show that in contrast to depletion of giantin, complete knockout of the gene does not prevent cells from generating cilia on serum withdrawal. We hypothesised that compensatory mechanisms might enable KO cells to produce normal cilia, while acute suppression of expression using RNAi did not allow such compensation. Such mechanisms would also explain the relatively mild manifestation of ciliary defects in giantin mutant zebrafish compared to *in vivo* morpholino knockdown. RNAseq showed that RCAN2 (Regulator of Calcineurin-2, also called calcipressin-2 and ZAKI-4) is the most strongly upregulated gene in these giantin KO cells. Here we show that RCAN2 localises to centrioles and its depletion affects ciliogenesis. Furthermore, removal of RCAN2 in giantin KO cells recapitulates the giantin RNAi ciliary length phenotype.

## Results and Discussion

### Giantin KO cells show no gross defects in ciliogenesis

Using RNAi, we have shown previously that a reduction in expression of giantin is associated with a ciliogenesis defect in cells (Asante et al., 2013). Giantin-depleted cells produce fewer cilia and those that remain are longer suggesting a defect in length control. In contrast, analysis of giantin knockout RPE-1 cells (Stevenson et al., 2017) shows no gross defects in ciliogenesis at the level of light microscopy (Fig. 1A, quantified in B) or any significant change in cilia length (Fig. 1A, quantified fully in later experiments). Using serial section transmission electron microscopy, we detect no obvious abnormalities in the structure of the basal body, appendages, or ciliary pocket (Fig. 1C, enlarged in Cii). Furthermore, Golgi structure appears normal (see (Stevenson et al., 2017)). This phenotypic discrepancy between knockdown and KO cells is consistent with our observations in zebrafish (Bergen et al., 2017) and led us to hypothesise that altered expression of another gene or genes compensates for loss of giantin. A caveat to these experiments is that we were only able to derive one knockout line for giantin in RPE1 cells.

**Figure 1:**
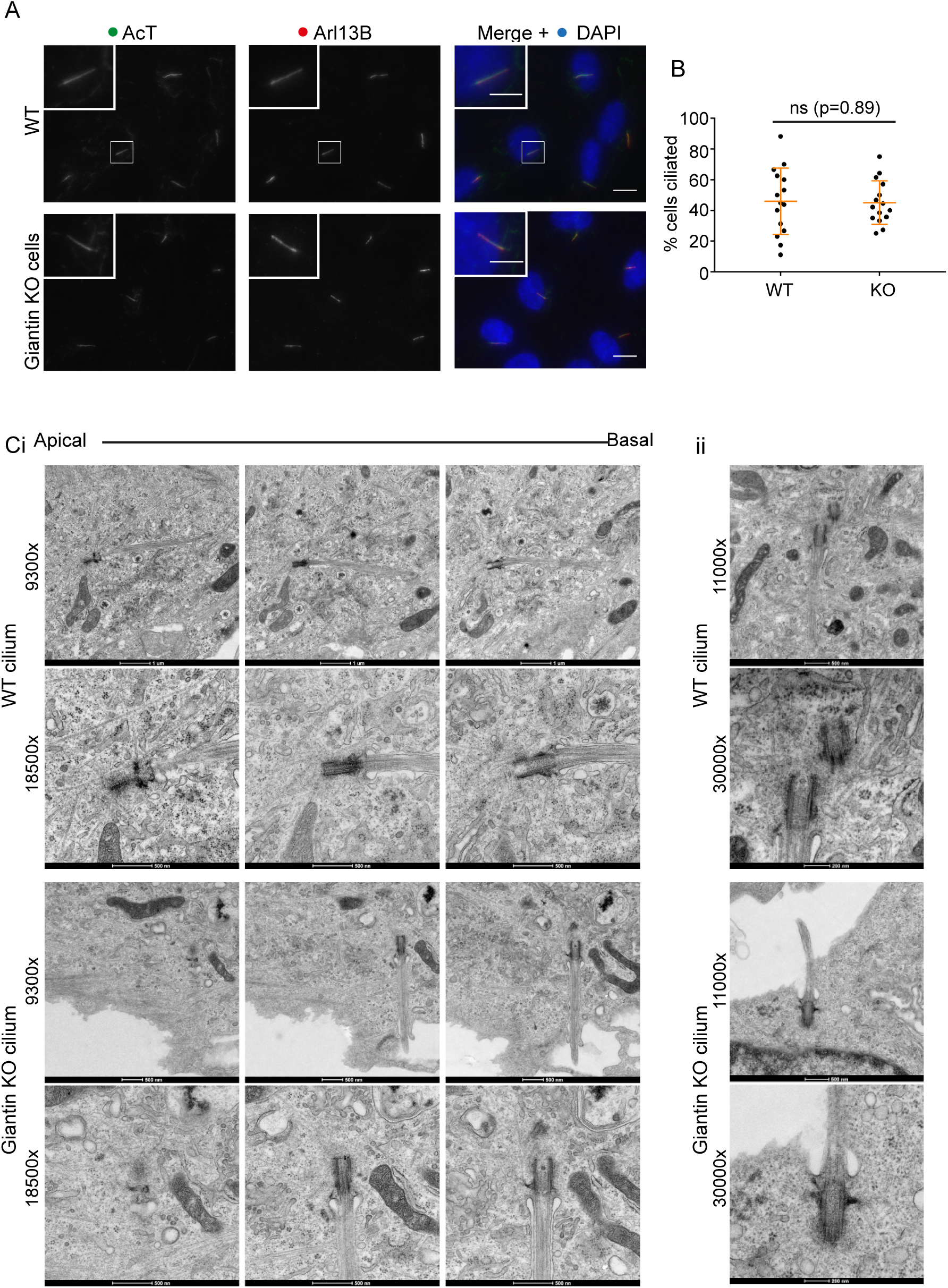
Giantin knockout RPE1 cells have no obvious defects in ciliogenesis. (A) Representative images of RPE1 WT and giantin KO cells fixed after 24 h of serum starvation and labelled for Arl13B (red), acetylated tubulin (green) and DAPI (blue). Scale bars are 10μm and 5μm on the merge and insert respectively. (B) Quantification of the experiment represented in A (n=3). The mean percentage of cells producing cilia does not change upon KO of giantin. Error bars represent SD, statistics Mann-Whitney. (C) TEM images of (i) serial and (ii) single plane 70 nm sections through representative WT and giantin KO cilia at two different magnifications.

RNAseq of triplicate RNA samples from these and control cells followed by pairwise statistical comparison identified 1025 genes whose expression level is decreased >4-fold. These data are described in Stevenson et al. (2017) and the raw data are available via the ArrayExpress database (www.ebi.ac.uk/arrayexpress) under accession number E-MTAB-5618. The most highly upregulated gene, and therefore the most likely to underpin any compensation for loss of giantin, is RCAN2. RCAN2 is a negative regulator of calcineurin and modulates Ca^2+^-dependent signalling. Calcineurin is required for transcriptional signalling; RCAN2 acts as a negative regulator of nuclear factor of activated T-cells (NFAT) activation and therefore of NFAT-dependent transcription. RNAseq showed that in giantin KO cells, RCAN2 is >256x upregulated compared to the parental cell line (Stevenson et al., 2017), implicating RCAN2 in adapting cells to the lack of giantin.

To determine whether RCAN2 upregulation is a conserved response to loss of giantin, we examined expression of RCAN2 in two giantin knockout (KO) zebrafish lines (Bergen et al., 2017). Using cDNA derived from young adult dissected jaw and skull bone and cartilage elements, qPCR expression analyses showed that zebrafish rcan2 is 3.9 and 2.5-fold up-regulated in giantin *X3078* and *Q2948X* mutant lines respectively, when normalised to β-actin (actb1) as control (Fig. 2A). Expression levels of hypoxanthine-guanine phosphoribosyl transferase 1 (hprt1) and glyceraldehyde-3-phosphate dehydrogenase (gapdh) were not significantly changed. This shows that upregulation of RCAN2 following loss of giantin is conserved and is therefore consistent with the hypothesis that the limited phenotypes seen in both giantin KO models are due to compensation. This also reduces concerns arising from the analysis of only one giantin knockout RPE1 cell line. We postulated that if this dramatic increase in expression is responsible for the limited phenotypes of giantin knockouts, then RCAN2 should have a key role in ciliogenesis.

**Figure 2:**
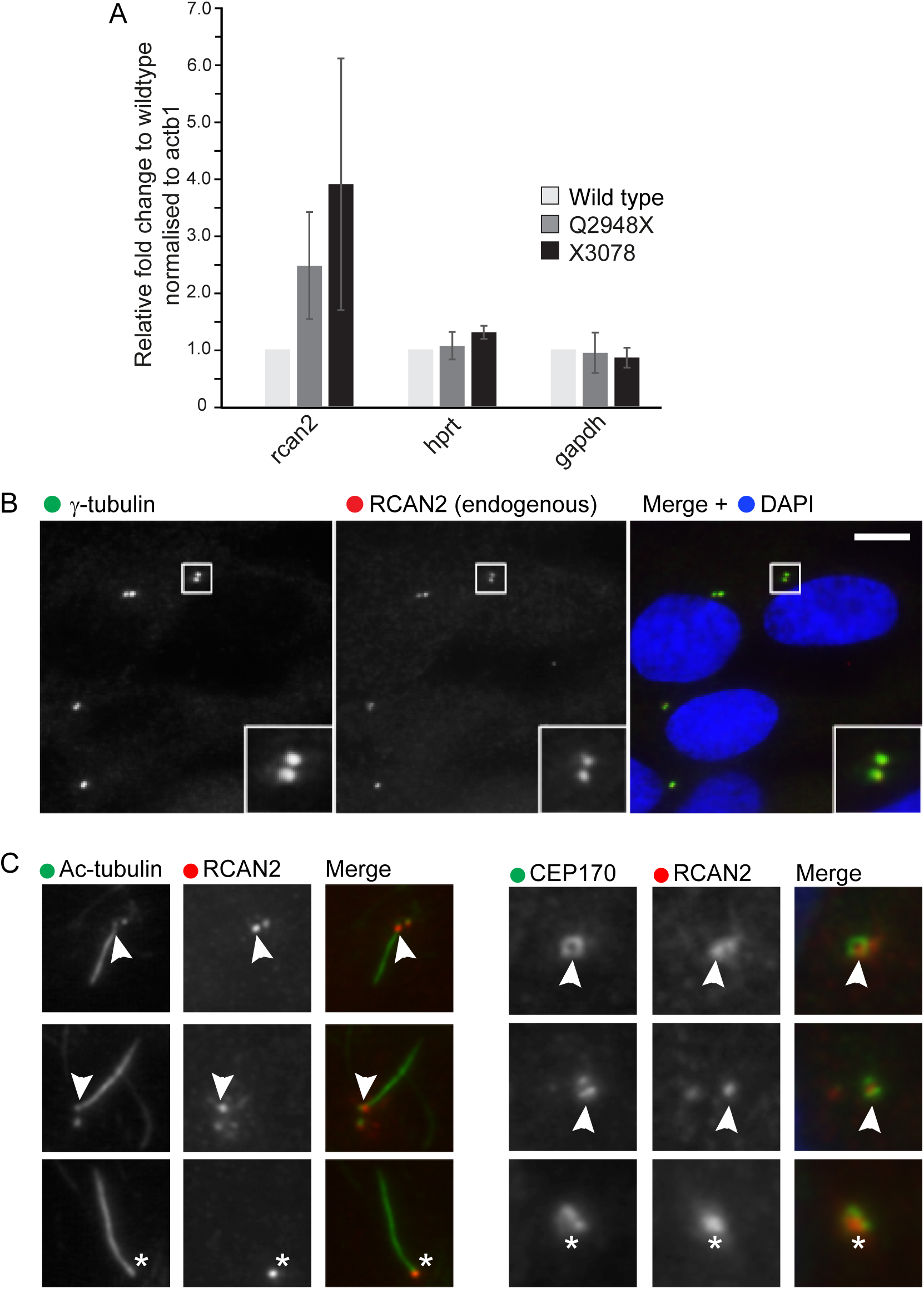
RCAN2 localises to centrioles. (A) QPCR of two giantin knockout zebrafish lines (Q2948X and X3078) compared to wild-type sibling controls shows that rcan2 is strongly upregulated when compared to the housekeeping gene beta-actin. In contrast, hprt1 and gapdh are not upregulated. (B) Endogenous RCAN2 (red) localises to centrioles (labelled with γ-tubulin (green) in hTERT-RPE1 cells. Bar =10 μm. (C) RCAN2 is frequently (25% +/-11%, n=3 independent experiments) found concentrated at the mother centriole (from which the cilium extends) as shown by acetylated tubulin labelling of the ciliary axoneme (arrowhead). Where only one puncta of RCAN2 labelling is found (asterisk, (7% of cells +/-3%, n=3 independent experiments)), a ciliary axoneme is always found extending from this centriole. Bar in B = 5 μm. (D) The distal appendage protein CEP170 is found associated with the brighter of the two RCAN2-positive centrioles (arrowhead) as well as with single centrioles positive for CEP170 (asterisk). Boxes are 5 × 5 μm.

The localisation of RCAN2 has not been reported. Immunofluorescence of RCAN2 in RPE-1 cells showed that it localised to centrioles (Fig. 2B) and occasionally showed enhanced localisation to the mother centriole (25% +/-11%, n=3 independent experiments) from which the primary cilium extends (shown by acetylated tubulin labelling in Fig. 2C). This was confirmed by labelling with the distal appendage protein CEP170 (Fig. 2D) which can be used as a marker of the mother centriole (Huang et al., 2017). In those cases where only one centriole is evident from RCAN2 labelling (7% of cells +/-3% n=3 independent experiments), we always see the axoneme extending from, and CEP170 associated with, that centriole.

To determine whether RCAN2 was compensating for giantin depletion with respect to cilium length, we depleted RCAN2 using two individual siRNA duplexes (Fig. 3) as well as a pool of 4 duplexes (Fig. S1). This experiment validated the specificity of the antibody used since centriolar labelling was lost following RNAi suppression of RCAN2 expression (Fig. 3A). The efficacy of suppression was measured as a function of the intensity of RCAN2 labelling at the centrioles (Fig 3B). Unfortunately, the antibody was not found suitable for immunoblotting. Depletion of RCAN2 did not affect the proportion of cells that could generate cilia following serum withdrawal (Fig. 3C). Quantification of these data showed that in WT RPE-1 cells, suppression of RCAN2 resulted in shorter cilia (Fig. 3D). In contrast, depletion of RCAN2 in giantin KO cells resulted in longer cilia, even when compared to controls. Our hypothesis was that RCAN2 could be suppressing the phenotypes seen following acute loss of giantin in RNAi experiments, namely an increase in cilia length. Consistent with this, we see that reducing the levels of RCAN2 in giantin KO cells recapitulates the longer cilia phenotypes seen on giantin knockdown in WT cells (Asante et al., 2013). Depletion of RCAN2 has no effect on the integrity of the Golgi, nor on targeting of the glycosyltransferase galactosyltransferase T (GalT) (Fig. S1B versus S1A). The efficacy of depletion of RCAN2 using a pool of 4 siRNA duplexes is also shown (Fig. S1D). It was not possible to study the effects of RCAN2 overexpression in these experiments as transfection with myc-RCAN2 blocked ciliogenesis in >80% of transfected cells. Attempts to titrate expression levels were unsuccessful.

**Figure 3:**
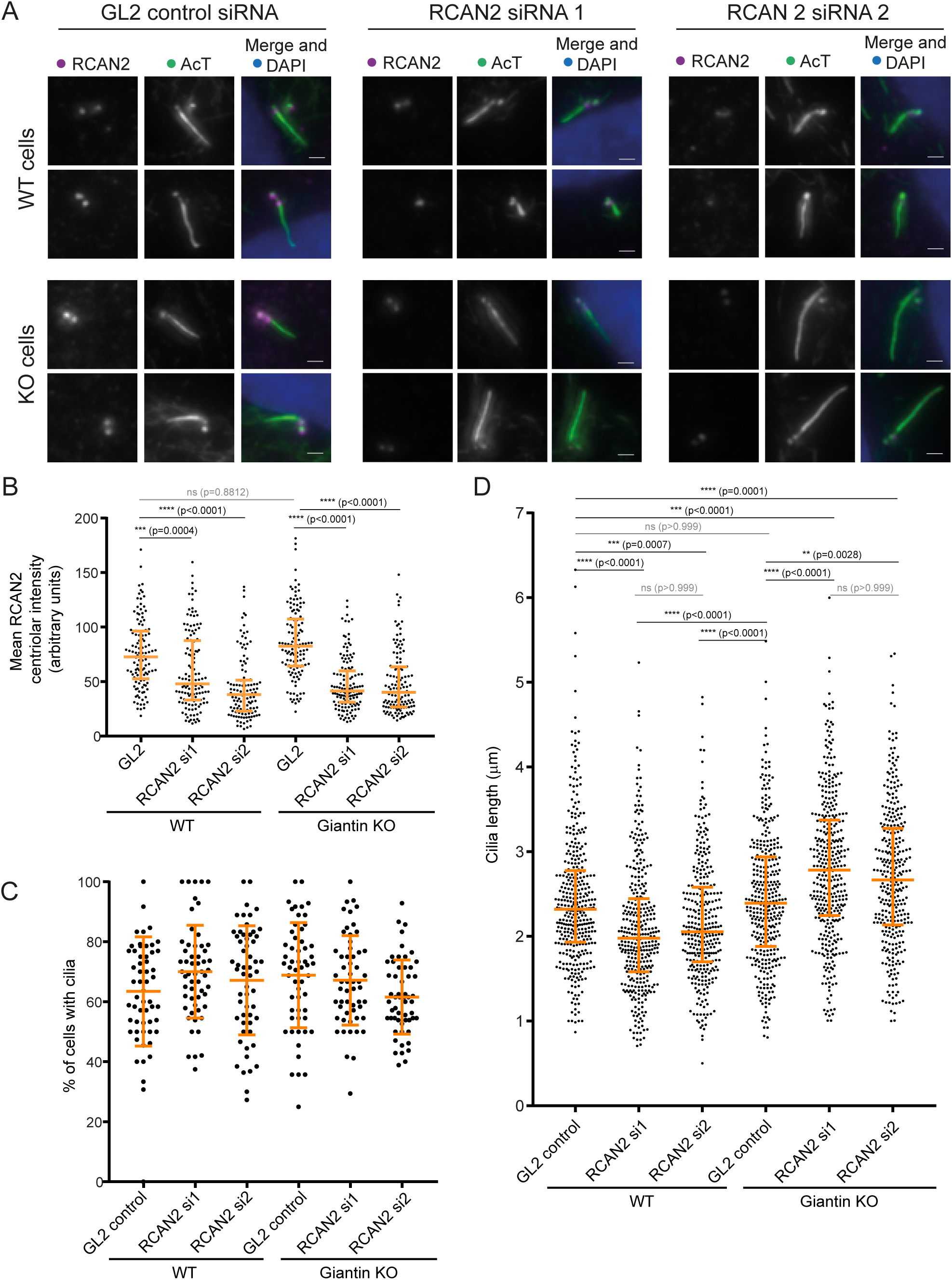
RCAN2 compensates for loss-of-function of giantin in cilia length control. (A) Depletion of RCAN2 using two different siRNA duplexes (1 and 2) is effective as judged by reduced centriolar labelling compared to cells transfected with control siRNA (GL2), quantified in (B). Depletion of RCAN2 results in shorter cilia in WT RPE-1 cells but longer cilia in giantin KO cells compared to GL2-transfected controls. Two examples are shown in each case. (C) Depletion of RCAN2 does not affect the ability of cells to produce cilia. (D) Quantitation of cilia length in RCAN2-depleted WT and giantin KO RPE-1 cells. Depletion of RCAN2 in wild-type RPE-1 cells results in a decrease in cilium length. In contrast, depletion of RCAN2 in giantin knockout RPE-1 cells increases cilia length, even when compared to WT cells. Statistical significance was tested using one-way, non-parametric ANOVA (Kruskal-Wallis test) with Dunn’s multiple comparisons test. >35 cilia were measured in each case from 3 independent biological replicates.

Our data strongly suggest that upregulation of RCAN2 ameliorates defects in cilia length control. The phenotypes we observe following acute and partial depletion of giantin in mammalian cells (Asante et al., 2013) and zebrafish embryos (Bergen et al., 2017) are not seen following permanent loss of function of giantin. The fact that RCAN2 depletion leads to shorter cilia in WT cells and longer cilia in giantin KO cells likely reflects a general loss of length control rather than a linear relationship whereby increasing RCAN2 levels lead to reducing cilium lengths. In WT cells depleted of RCAN2, cilia are unable to maintain their length and so become shorter, whereas in giantin KO cells the loss of length regulation means the effects of giantin loss become dominant and cilia become longer.

It is interesting that our data implicate cilium length control by compensatory pathways *in vitro* in proliferating cells. The importance of cilia to cell growth in culture is poorly understood and so we cannot predict whether defective ciliogenesis alone would provide sufficient adaptive pressure for RCAN2 upregulation. It is possible however that loss of giantin results in a pro-ciliary phenotype which is compensated by upregulation of RCAN2. In support of this, we find that overexpression of myc-RCAN2 results in far fewer cells (<20%) that produce cilia (Fig. 4A, n=100 cells) and indeed incidences (~10% of myc-RCAN2-expressing cells) where we can detect Arl13b positive structures that are devoid of acetylated-tubulin labelling (Fig. 4B). These are similar in appearance to “decapitated” cilia that have been described by others to drive cilia resorption and cell cycle progression (Phua et al., 2017). In addition, RCAN2 may be acting as a regulator of gene transcription through NFAT signalling to correct other deficiencies in the KO cells. For example, we have shown previously that the glycosyltransferase content of the Golgi apparatus is altered in giantin KO cells suggesting glycosylation defects arising from loss of giantin function might have been corrected to generate bioequivalent glycans (Stevenson et al 2017).

Many studies have shown that RCAN2 regulates calcineurin-dependent activation of NFAT signalling and subsequently NFAT-dependent transcriptional programmes. To confirm that upregulation of RCAN2 in giantin KO cells is of functional importance, we examined NFAT localisation in WT and giantin KO cells by immunofluorescence (Fig. 4C). As shown in Fig. 4D, the ratio of nuclear vs cytoplasmic levels of NFAT1 was higher in giantin KO cells compared to WT. This nuclear localisation was further enhanced in response to serum starvation. The distribution of NFAT1 in cells depleted of giantin by shRNA was similar to that in wild-type cells, supporting our argument that acute giantin depletion does not induce adaptation in these timescales, hence the longer cilia. The higher RCAN2 expression level seen in the giantin KO cells thus could result in greater activation of the calcineurin/NFAT signalling pathways. Although RCAN2 is commonly considered to be a negative regulator of calcineurin (Cao et al., 2002; Loh et al., 1996) other studies have found it can positively promote NFAT signalling (Sanna et al., 2006) as appears to be the case here. Alternatively, enhanced RCAN2 expression could result from attempts to suppress upregulated calcineurin-based signalling. Activation of NFAT signalling has been described to occur downstream of α6β4 integrin activation in metastatic cancer cells (Jauliac et al., 2002) and RCAN1 has been shown to modulate cancer cell migration (Espinosa et al., 2009). Of note, both giantin KO cells and giantin KO fish have substantial defects in extracellular matrix related functions (Bergen et al., 2017, Stevenson et al., 2017). Thus, we speculate that changes in matrix glycoproteins could be the trigger for compensatory changes in gene expression in giantin knockout models.

Rodent knockouts for giantin (rat (Katayama et al., 2011) and mouse (Lan et al., 2016)) exhibit defects in extracellular matrix secretion and assembly, cartilage and bone formation, and have syndromic cleft palate. RCAN2 KO mice are viable but show selective defects in osteoblast function (Bassett et al., 2012), in impaired intramembranous ossification, and in cortical bone formation. Indeed, loss of RCAN2 function is associated with reduced bone mass whereas giantin mutant zebrafish (with increased RCAN2) shows enhanced bone formation in the intervertebral discs (Stevenson et al., 2017). Evidence also exists that inhibition of calcineurin using cyclosporin is linked to cilia function, shown by the presence of abnormal basal bodies in proximal convoluted tubule cells from patients receiving Cyclosporin A (Kirwan, 1982). Ciliary dysfunction is frequently linked with skeletal defects. Indeed, the “skeletal ciliopathies” are grouped according to phenotypic descriptions of skeletal defects (Huber and Cormier-Daire, 2012). These diseases include Jeune syndrome in which subunits of the dynein-2 motor are mutated. Intriguingly, our work has led from the biology of giantin to the dynein-2 IFT motor (Asante et al., 2013) to the identification of RCAN2 as a key regulator of ciliary function.

These and other findings implicate RCAN2 and calcineurin in transcriptional regulation via cilia. This provides a potential point of integration at the nexus of Ca^2+^ and ciliary signalling. In support of this, tax-6, the *C. elegans* orthologue of calcineurin, regulates the trafficking of the key ciliary protein, polycystin-2 (PKD2) (Hu et al., 2006). PKD2 is a Ca^2+^ permeable cation channel and is mutated in 15% of cases of autosomal dominant polycystic kidney disease. Of note here, the partner of PKD2, PKD1 (mutated in the other 85% of ADPKD cases) is understood in osteoblasts to be a mechanoreceptor transducing signals from the cilia to induce osteoblastic gene transcription by potentiating Ca^2+^-dependent calcineurin/NFAT signalling (Dalagiorgou et al., 2013). Its role in canonical NFAT signalling suggests a mechanism by which it could regulate coupling of ciliary Ca^2+^ to cellular functions through transcriptional control. Our findings also have significant potential for understanding the mechanisms of cilia-based Ca^2+^-dependent signal transduction, including modulation of Shh signalling through Gli (Delling et al., 2013). One possibility is that the concentration of RCAN2 at the base of the cilium acts to quench calcium-dependent calcineurin signalling restricting it to the cilium itself. Understanding the role of RCAN2 in ciliary signalling will therefore have significant implications for diverse areas of fundamental importance to our understanding of cell function as well as of clinical relevance including bone formation, renal function, developmental signalling and mechanosensing.

**Figure 4:**
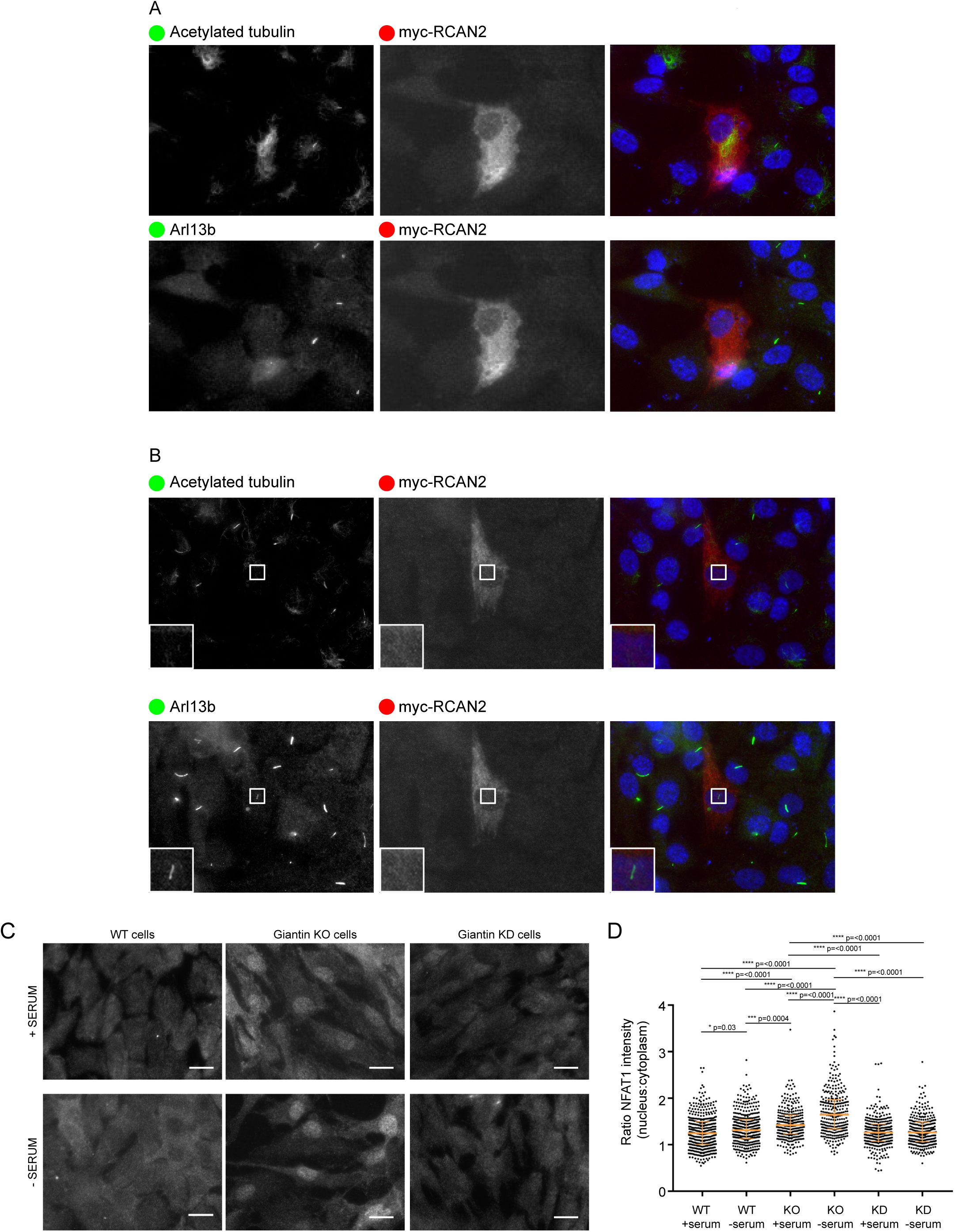
Giantin KO cells show higher levels of NFAT1 activation. (A) Expression of myc-RCAN2 (green in merge) prevents cilia formation as shown by acetylated tubulin or Arl13b (both pseudocoloured red in merge; images were obtained from triple-labelled cells with Alexa-488, −568 and −647). 80% ofmyc-RCAN2-expressing cells failed to extend a cilium (n=100). (B) In 5% of these 100 myc-RCAN2-expressing cells, we observed “decapitated” cilia, positive for Arl13b but not for acetylated tubulin. (C) Representative images showing WT and giantin KO and knockdown (KD) cells grown in serum or serum deprived for 24 hours and then immunolabelled for NFAT1. Scale bar 10 μm. (D) Quantification of the ratio of nuclear: cytoplasmic NFAT1 staining intensity in experiments represented in A (n=3, bars represent median and interquartile range, p values calculated using non-parametric ANOVA (Kruskal-Wallis test) with Dunn’s multiple comparisons test).

## Author contributions

We would like to thank Roderick Skinner for his help with zebrafish related work and the Wolfson Bioimaging Facility staff for help with microscopy. NLS and DJB designed and performed experiments and analysed data. AX, EW, FH, JM, and LV performed experiments and analysed data, DJS and CLH conceived and managed the project and contributed to data analysis. All authors co-wrote the manuscript.

## Acknowledgements

We would like to thank the Earlham Institute for the RNAseq analysis. We also thank the MRC and Wolfson Foundation for establishing the Wolfson Bioimaging Facility, and confocal microscopy was supported by a BBSRC ALERT 13 capital grant (BB/L014181/1). The project was funded by the MRC (MR/K018019/1), the Wellcome Trust (099848/Z/12/Z), and the University of Bristol.

## Materials and Methods

All reagents were purchased from Sigma-Aldrich unless stated otherwise.

### Cell culture

Human telomerase-immortalised retinal pigment epithelial cells (hTERT-RPE-1, ATCC) were grown in DMEM-F12 supplemented with 10% FCS (Life Technologies, Paisley, UK). Cell lines were not authenticated after purchase other than confirming absence of mycoplasma contamination. Transfections were performed using Lipofectamine 2000™ according to the manufacturer’s instructions (Invitrogen, Carlsbad, CA). Myc-DDK-tagged-human regulator of calcineurin 2 (RCAN2, accession number NM_005822.2) was purchased from Generon (Maidenhead, UK).

### Knockout models

Giantin KO RPE-1 cells are described in (Stevenson et al., 2017). Giantin KO zebrafish are described in (Bergen et al., 2017). London AB, Tupfel long fin (TL), and AB/TL zebrafish were used and maintained according to standard conditions (Westerfield, 2000), kept in 3.5 litre tanks in groups of max 15 adult fish, and staged accordingly (Kimmel et al., 1995). Ethical approval was obtained from the University of Bristol Ethical Review Committee using the Home Office Project License number 30/2863.

### RNAi

RCAN2 was depleted in RPE-1 cells using RNA interference. siRNA duplexes directed against RCAN2 were made by Dharmacon (MQ-020054-01-0002 siGENOME Human RCAN2 (10231), Dharmacon, GE Healthcare). Transfections were carried out using either a control or RCAN2 siRNA. Four pooled siRNAs were used to deplete RCAN2, with the following target sequences, RCAN2 siRNA #1 (GGAUAGAGCUUCAUGAAAC), RCAN2 siRNA #2 (CGUAUAAACUUCAGCAAUC), RCAN2 siRNA #3 (CCACGCCAGUCCUCAACUA) and RCAN2 siRNA #4 (GCUCUACUUUGCACAGGUU). GL2 control siRNA (CGUACGCGGAAUACUUCGAUU) was used as a negative control. All siRNA reagents were prepared in 2M CaCl_2_and incubated for 5 min at room temperature before an equal volume of 2X BES buffered solution (BBS) was added to the solution and allowed to equilibrate for a further 30 min at room temperature. Final siRNA concentrations used were 5 μM. Confluent RPE-1 cells grown on round 22 mm glass coverslips in Costar 6-well plates. Cells were incubated at 37°C, 3% CO2 for 24 h before replacing the growth media. Cells were grown for an additional 24 h at 37°C, 5% CO2 then serum starved for ciliogenesis assays.

### Antibodies, labelling and microscopy

Antibodies used: mouse monoclonal anti-giantin (full length, Abcam, Cambridge, UK, ab37266), rabbit polyclonal anti-giantin (N-terminus, Covance, CA, PRB-114C), rabbit anti-RCAN2 (GTX31373, Insight Biotechnology, London UK), anti-Arl13b (17711-1-AP, Proteintech, Manchester, UK), sheep anti-my was a kind gift from Harry Mellor (University of Bristol), sheep anti-GRASP65 was a kind gift from Jon Lane (University of Bristol), anti-GalT (CB002, CellMab, Gothenberg, SE), anti-NFAT1 (GTX127932, GeneTex, Insight Biotechnology, London UK).

For antibody labelling, cells were grown on autoclaved coverslips (Menzel #1.5, Fisher Scientific, Loughborough, UK), rinsed with PBS and fixed in MeOH for 4 minutes at −20°C. Cells were then blocked in 3% BSA-PBS for 30 minutes and incubated with primary then secondary antibody for 1 hour each, washing in between. Nuclei were stained with DAPI [4,6-diamidino-2-phenylindole (Life Technologies, Paisley, UK, D1306)] for 3 minutes and coverslips mounted in Mowiol (MSD, Hertfordshire, UK) or Prolong Diamond antifade (Thermo Fisher, Paisley, UK). Fixed cells were imaged using an Olympus IX70 microscope with 60x 1.42 NA oil-immersion lens, Exfo 120 metal halide illumination with excitation, dichroic and emission filters (Semrock, Rochester, NY), and a Photometrics Coolsnap HQ2 CCD, controlled by Volocity 5.4.1 (Perkin Elmer, Seer Green, UK). Chromatic shifts in images were registration corrected using TetraSpek fluorescent beads (Thermo Fisher). Images were acquired as 0.2µm z-stacks.

### Reverse transcriptase PCR and quantitative Real-Time PCR

Young adult fish (60 and 63 days post fertilisation for Q2948X (F4) and X3078 (F3) lines respectively, 3 individuals each genotype group) were lethally anaesthetised (<0.15% Tricaine) and their frontal skull and ventral jaw bone and cartilage elements were carefully dissected using a standard dissecting microscope. From these separate samples, total RNA was isolated using RNeasy mini kit (cat# 74104, Qiagen) and reverse transcriptase reaction was performed by using Superscript III (cat# 18080093, ThermoFisher Scientific) according to the manufacturers’ protocols. Quantitative Real-Time PCR (qPCR) reaction [Primers: *rcan2* (ENSDART00000143379.2): F-5’CCAGGTGCAGAACCCAGTAT; R-5’CCTCAGACCCTGGTTGTGTT, *hprt1:* F-5’ATGGACCGAACTGAACGTCT; R-5’GGTCTGTATCCAACGCTCCT, *actb1*: F-5’TCAACACCCCTGCCATGTAT, R-5’CAGGAAGGAAGGCTGGAAGA; *gapdh*: F-5’TGTTCCAGTACGACTCCACC, R-5’GCCATACCAGTAAGCTTGCC] was undertaken with DyNAmo HS SYBR green (F410L, ThermoFisher Scientific) on 175 ng/µl cDNA libraries, cycling (40 times) at 95°C 25 seconds, 57.5°C 30 seconds, and 70°C 45 seconds followed by a standard melt curve (QuantStudio3, Applied Biosystems).

### RNAseq

Triplicate samples of total RNA (three independent passages) from WT and giantin KO RPE-1 cells were analysed by RNAseq by the Earlham Institute (formerly The Genome Analysis Centre). The methodology and analysis is fully described in (Stevenson et al., 2017).

### Quantification and statistical analysis

Statistical analyses were performed using Graphpad Prism 7.00. The tests used and sample sizes are indicated in the figure legends, p-values are shown on the figures. All tests met standard assumptions and the variation between each group is shown. Sample sizes were chosen based on previous, similar experimental outcomes and based on standard assumptions. No samples were excluded. Randomisation and blinding were not used except where the genotype of zebrafish was determined after experimentation. Statistical significance was tested using one-way, non-parametric ANOVA (Kruskal-Wallis test) with multiple comparisons using Dunn’s test.

### Data and software availability

Raw RNAseq data are available in the ArrayExpress database (www.ebi.ac.uk/arrayexpress) under accession number E-MTAB-5618.

**Figure S1:**
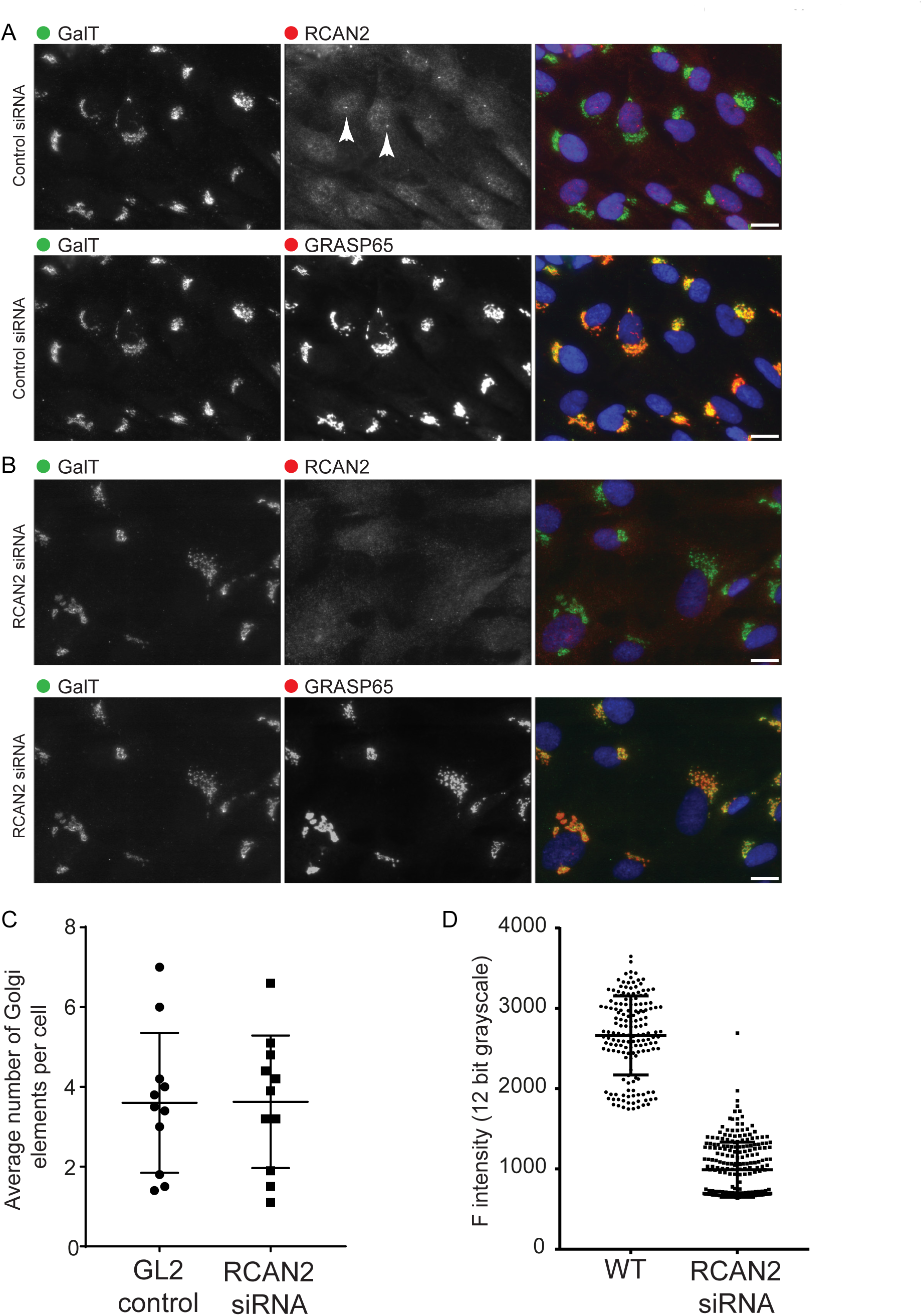
RCAN2 is not required to maintain Golgi structure. (A, B) Depletion of RCAN2 in RPE-1 cells (B) does not affect targeting of GalT (galactosyltransferase) to the Golgi compared to controls (A). Cells were labelled with antibodies to detect GalT (green), RCAN2 (red, top panels) and GRASP65 (far red, bottom panels). The localisation of RCAN2 to centrioles (arrowheads) is lost following depletion using RNAi. Golgi structure is not grossly affected by depletion of RCAN2. Images shown are from WT RPE-1 cells, the same effect is seen when using giantin KO cells. Bar = 10 μm. (C) The average number of Golgi elements per cells (dots show the means of 10 fields of view) does not differ between control and RCAN2-depleted cells. (D) Quantification of the intensity of RCAN2 labelling shows the efficacy of the siRNA in these specific experiments, >50 cells in each of n=3 independent replicates.

